# Evolution of linkage and genome expansion in protocells

**DOI:** 10.1101/746495

**Authors:** András Szilágyi, Viktor Péter Kovács, Eörs Szathmáry, Mauro Santos

**Affiliations:** Department of Plant Systematics, Ecology and Theoretical Biology, Eötvös Loránd University, 1117 Budapest, Hungary; Evolutionary Systems Research Group, Centre for Ecological Research, Hungarian Academy of Sciences, 8237 Tihany, Hungary; Center for the Conceptual Foundations of Science, Parmenides Foundation, 82049 Pullach/Munich, Germany; Grup de Genòmica, Bioinformàtica I Biologia Evolutiva (GGBE), Departament de Genètica i de Microbiologia, Universitat Autonòma de Barcelona, Bellaterra, Barcelona, Spain

**Keywords:** Chromosomes, intragenomic conflict, parasites, antagonistic epistasis, stochastic corrector model

## Abstract

Chromosomes are likely to have followed unlinked genes in early evolution. Genetic linkage reduces the assortment load and intragenomic conflict in reproducing protocell models to the extent that chromosomes can go to fixation even if chromosomes suffer from a replicative disadvantage, relative to unlinked genes, proportional to their length. Here we show that chromosomes spread within protocells even if recurrent deleterious mutations affecting replicating genes (as ribozymes) are taken into account. Dosage effect selects for optimal genomic composition within protocells that carries over to the genic composition of emerging chromosomes. Lacking an accurate segregation mechanism protocells continue to benefit from the stochastic corrector principle (group selection of early replicators), but now at the chromosome level. A remarkable feature of this process is the appearance of multigene families (in optimal genic proportions) on chromosomes. An added benefit of chromosome formation is an increase in the selectively maintainable genome size (number of different genes), primarily due to the marked reduction of the assortment load. This result complements the established benefit conferred by chromosomes on protocells allowing for the fixation of highly specific and efficient enzymes.

## Introduction

No present living organism can come about without the replication of its entire genetic information contained in the chromosomes. Furthermore, chromosomes are a prerequisite for the evolution of complex metabolism through the appearance of specific enzymes (1). How did chromosomes originate in the first place? The primeval self-replicating entities were probably naked RNA molecules coexisting as surface-bound populations that had to meet some stringent criteria were they able to evolve toward higher-level units of selection such as protocells; namely, the entities enclosing functional replicators (molecules serving as both templates and catalysts) into amphiphilic vesicles (2,3,4). Protocells alleviate some obstacles faced by prebiotic systems as they increase interactions among hosted molecules and confer robustness against parasitic replicators through group selection (5,6,7).

However, because the genetic information within ancient protocells was likely segmented (2), unlinked replicators competed among themselves for shared resources because their relative growth rates were not under the control of the protocell. This imposed a first level of selection due to the internal competition of replicators that functioned for their own good (8). Offspring protocells could have inherited an unbalanced set of genes, be unable to grow and reproduce (assortment load). Clonal selection guaranteed that those protocell lineages hosting cooperative genes would proliferate and eventually take over (5). Although the stochastic assortment effects vanish with increasing redundancy of each sequence type, this is an unrealistic scenario for at least two reasons. First, with high redundancy there is the risk that Darwinian selection would be stopped because of dilution of favorable mutations (9). Second, high redundancy increases the mutational load and eventually pushes the population towards extinction (10). Furthermore, notwithstanding some claims on the putatively large number of different gene types that could be hosted by protocells (11), recent experiments have shown that the number of independent templates per protocell must be sufficiently small for protocells to be evolutionarily stable (12). At some point in time the linkage of genes in one continuous chromosome occurred (6,13), but it is still unclear how this could have happened.

Previous attempts to explain the origin of chromosome were limited in their scope because they modelled some very specific scenarios: only two genes, no dosage effect and absence of deleterious mutations (14), which are known to place severe limitations to the upper bound of informational length because of the error-catastrophe problem; that is, when the amount of information lost through the continuous input of deleterious mutations (mutation load) is higher than the amount of information that natural selection can recover (15,16). The hurdles of assortment and mutation genetic loads faced by protocells should have been related problems concerning selection for linkage. Extensive theoretical work has shown that epistasis (understood as the departure from multiplicative selection) is critically important for the evolution of linkage (17,18): when loci are subjected to recurrent deleterious mutations, linkage is always favored with positive epistasis (i.e., when mutations have a weaker effects on fitness when combined). Therefore, if positive epistasis was common in early genetic systems we might expect that there was strong selection for linkage (chromosomes) because this would simultaneously reduce the two types of genetic load.

Using metabolic control theory, Szathmáry (19) showed that for a linear metabolic pathway deleterious mutations that affect different enzymes in the pathway exhibit positive epistasis when selection is for maximum flux. Starvation is a common condition in present day bacteria (20) and probably was so in early protocells, which suggests that protocell’s fitness was mainly determined by the flux of a non-saturated pathway metabolizing limiting nutrients. Here we show that the major evolutionary transition “independent replicators → chromosomes” (13) was strongly favorable in early protocells and opened new routes to the evolution of complexity.

## Model

Our goal is to understand the evolution of chromosomes and genome expansion from first principles. A key ingredient is the suggestion (21) that primordial ribogenes were replicated in a manner similar to present-day Q*β* phage RNA, with tRNA-like 3′ tags (i.e., a recognition site for the replicase at the end of the template; see ref. 22). Therefore, the genes within protocells are organized as having a target region of *η*_*t*_ = 20 nucleotides that defines an average affinity towards the replicase (i.e., whether they are good substrates for the replicase), plus a sequence of *η*_*m*_ = 80 nucleotides involved in their metabolic function. For simplicity, we assume that protocells hosted both replicase and ligase ribozymes in high concentration, and only follow the dynamics of the joining of templates to form longer polymers *G*_*k*_ + *G*_*l*_ → *G*_*k*_ ⋅ *G*_*l*_ (*k,l* = {1,…, *D*}), where *G*_*k*_ stands for a metabolic gene *k* essential for protocell survival (*k* and *l* are not necessarily different; i.e., the same gene can be present in multiple copies) and *D* is the number of essential genes. The ligase can act in a similar manner between chromosomes 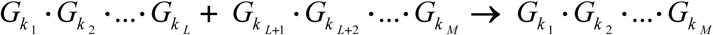, (*k*_1_,*k*_2_,…, *k*_*M*_ = {1,…, *D*}). Note that ligase (linking segments) and replicase (copying genes and chromosomes) ribozymes are not represented in the model as genes.

The population consists of *N* protocells at *t*_0_; all genes have maximum metabolic and replicative activity and each cell contains a random composition of the *D* essential genes. The total number of genes in a cell is a uniform random number between 1 and *S* − 1, where *S* is the maximum number of genes (independent from activity type) in a protocell. Protocell’s fitness is

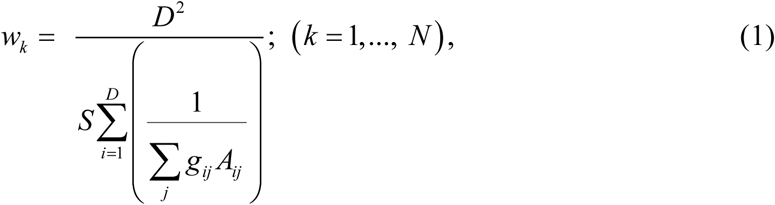

where *A*_*ij*_ and *g*_*ij*_ are the metabolic activity and copy number of the *j*th variant of gene *i*. Eq. (1) has its maximum if all enzymes have the same total activity (*SI Section 1*); and fitness is always reduced even in the presence of compensatory mutations (*SI Section 2*). Eq. (1) also captures the effect of the dosage.

Each type of metabolic activity corresponds to a specific nucleotide pattern of the metabolic region. In our binary (0, 1) representation, a sequence with a block of eight consecutive 1s between positions 10*i* −9 and 10*i*, and 0 values otherwise (*i* = 1,2,…,*D*). In this model the maximum number of different metabolic activities is 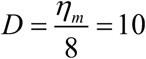. (This choice practically excludes enzymatic promiscuity.) The activity of a gene variant depends on the number of mutated bases according to the following

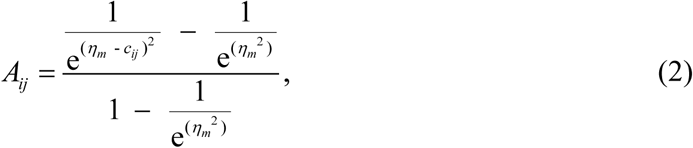

where *c*_*ij*_ stands for the number of non-mutated nucleotides in the metabolic region of gene version *j* of gene type *i* (23). Note that if *η*_*m*_ > 3, the activity simplifies to 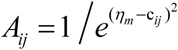, which depends on the number of mutated nucleotides only and does not reflect to the length. Eq. (1) relies on the assumption that fitness is – as usual for microbes – essentially determined by the flux of a linear pathway of reactions catalyzed by unsaturated enzymes. The fitness function is normalized; i.e., in the optimal case of no mutant copies and all essential enzymes present it is *S*/*D*. This function captures both the effect of deleterious mutations and the enzyme dosage (c.f. eq. 5 in ref. 24). (Note that if any essential gene is missing, the fitness of the protocell is zero.)

Within protocells, a template is replicated according to its replication probability, which depends on its target affinity towards the replicase

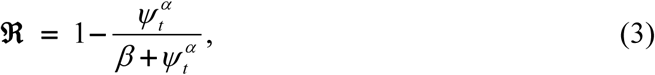

where *ψ*_*t*_ is the number of mutated nucleotides in the target region of the template, and *α*, *β* are constants. Eq. (3) decreases from a maximum of 1 with no deleterious mutations; in the following we use *α* = 5, *β* = 15 (25). During replication (only a single replicase acts on a template at a time) mutations are introduced at a rate *μ* per nucleotide, and the replicated template is added to the protocells’ set. We assume that the replicase travels at a constant speed along the template, which means that replication of a chromosome with *n* linked genes takes *n* times the time it takes to replicate a single gene (i.e., a *n*-fold selective disadvantage).

Regardless of whether or not a template is replicated, we assume that with probability *ν*_linkage_ two randomly chosen templates in the protocell will be linked into a longer template. The mechanism follows a restricted copy-choice (26): *i*) a replicase can switch from one template to another after copying a whole gene, the replicase stops after the second partner has been replicated; *ii*) two chromosomes can recombine if the gene type at the switching point is the same for both partners (e.g. ABE + DBCC → ABCC, and DBCC + ABE → DBE, etc.); *iii*) two genes can be linked to a chromosome without restriction; and *iv*) if the resulting chromosome is longer than a maximum limit (*MC*), there is no linkage (for computational reasons). Furthermore, a chromosome can break into two parts (between genes) with probability *ν*_break_. Besides linkage and break, we also implemented recombination between two random templates with probability *ν*_recomb_. The resulting template must contain at least one gene from both partners, and must be shorter than the limit *MC*. A protocell splits into two (by hypergeometric sampling, i.e. no replacement) when the number of genes reaches the maximum size *S*. The population dynamics follows a Moran (27) process where one daughter protocell replaces the parental protocell, and the other daughter protocell replaces a random member of the population. All simulations were performed in C; the source code is available upon request.

## Results and Discussion

### Positive (antagonistic or diminishing-returns) epistasis

As a consequence of the input of deleterious mutations in the different genes the direction of epistasis is positive, meaning that mutations have a weaker effect on protocell’s fitness when combined (*SI Section 3*). Under this condition, decreased recombination is always favored (17). It is worth mentioning that antagonistic epistasis has been predicted from studies of the effect of mutations on RNA folding (28) and analyses of RNA viruses (29), as well as in *E. coli* and *S. cerevisiae* using flux balance analysis and in silico studies of metabolic networks (30).

### Chromosomatisation and genome expansion

We first summarize the main findings and then focus on a particular scenario to understand the dynamics of the system.

In all analyzed situations, chromosomes always spread despite strong within-protocell selection against them. Even if a long chromosome breaks, a diverse set of smaller chromosomes with different number of genes can be present at equilibrium. However, in all cases chromosomes with full set(s) of genes dominate the system. If the split size *S* is low (i.e., if the maximal number of genes at the time of protocell division is low), chromosomes with one full set of essential genes are present in relatively high concentration. With increasing split size the concentration of chromosomes with two (or more) full sets of genes increases; that is, we observe a genome expansion of linked genes as a function of split size. Chromosome breakage produces solitary genes and shorter chromosomes that contain no full sets of genes, reducing the average length of chromosomes and protocell fitness. Nevertheless, in the transient period chromosome breakage introduces the necessary variation to reach an optimal composition of genes in the chromosome. Without chromosome breakage the system could freeze in a suboptimal state and, in equilibrium, only a few types of chromosomes remain in the system that excludes further optimization. Finally, we have found that recombination does not affect the results qualitatively.

We now focus on a particular case assuming *D* = 3 essential genes (A, B and C). We integrated the system until equilibrium, and at *t* = 10^6^ recorded the number of different types of genes in chromosomes with different gene numbers (Table 1). The most frequent chromosome (~ 50%) in the population was almost perfectly balanced with genes ABC, and the second most frequent (~ 21%) chromosome with genes ABCABC. Balanced ABCABCABC chromosomes (~ 1.5%) were also present. In other cases the gene composition was less balanced, but on the whole there is an almost perfect equilibrium in gene composition at the population level. Breaks produce solitary genes recurrently and because of the assortment load the ratio of solitary genes of different types is not well balanced.

**Table 1.** Number of different genes in chromosomes (and their ratio relative to the total) of different lengths (sum over the whole population). Parameters are: *D* = 3, *S* = 21, *μ* = 10^−3^, *ν*_linkage_ = *ν*_break_ = *ν*_recomb_ = 0.01.

| Number of genes<br>in chromosomes | Frequency | Number of<br>gene type A<br>(ratio) | Number of<br>gene type B<br>(ratio) | Number of<br>gene type C<br>(ratio) |
| --- | --- | --- | --- | --- |
| 1 | 0.0617 | 977 (0.2996) | 1010 (0.3097) | 1274 (0.3907) |
| 2 | 0.0796 | 1456 (0.3463) | 1510 (0.3591) | 1239 (0.2946) |
| 3 | 0.4939 | **8694 (0.3332)** | **8698 (0.3333)** | **8703 (0.3335)** |
| 4 | 0.0612 | 1113 (0.3439) | 991 (0.3062) | 1132 (0.3498) |
| 5 | 0.0415 | 798 (0.3642) | 764 (0.3487) | 629 (0.2871) |
| 6 | 0.2138 | **3768 (0.3335)** | **3771 (0.3338)** | **3759 (0.3327)** |
| 7 | 0.0057 | 92 (0.3077) | 109 (0.3645) | 98 (0.3278) |
| 8 | 0.0166 | 260 (0.2965) | 316 (0.3603) | 301 (0.3432) |
| 9 | 0.0152 | **261 (0.3242)** | **277 (0.3441)** | **267 (0.3317)** |
| 10 | 0.0108 | 181 (0.3175) | 203 (0.3561) | 186 (0.3263) |

Therefore, one of our main findings is that chromosomes with *n*⋅ *D* sets of genes can easily arise. This likely represented an important source of novelty in protocells’ evolution by allowing an expanded repertoire of metabolic activities through modification of existing genes (31,32) and, at the same time, without imposing an unbearable assortment load (1). In a sense, our findings exemplify Ohno’s (33; see also 34) model suggesting that duplication of parts of chromosomes is central to the evolution of new biological functions. The genome expansion of linked genes is most evident if we assume *ν*_break_ = 0, which also illustrates an important feature of the chromosomatisation dynamics (Fig. 1). Thus, because the formation of a (e.g.) 3−set balanced chromosome (ABC) has to overcome a strong within-protocell selection, what can be seen from Fig. 1 is that at the beginning 2-gene chromosomes increase in frequency at the expense of solitary genes; afterwards, 3-gene (balanced) chromosomes start increasing in frequency at the expense of 2-gene chromosomes; etc. In other words, the formation of chromosomes with *n*⋅ *D* sets of genes happens in a sort of stepwise process, which helps lessening the strong within-protocell selection against chromosomatisation. All imbalanced chromosomes are selected against, thus in equilibrium only chromosomes consisting of 3*n* genes are present and other gene numbers are unreachable by the system.

**Fig. 1.**
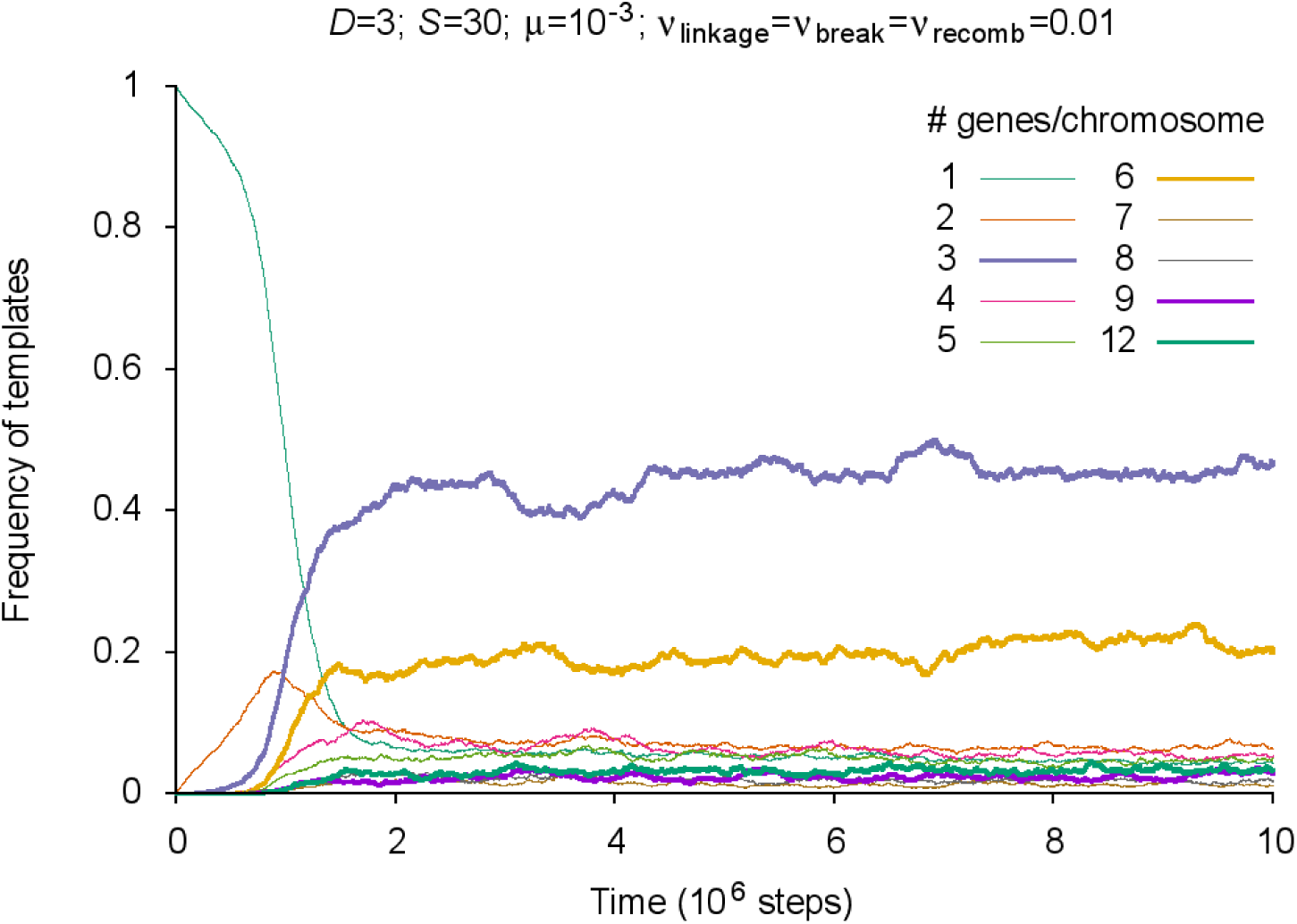
Semi-log plot of the frequency of different templates normalized on gene count (i.e., a chromosome with 3 genes counts as three when measuring the frequency) with no chromosome breakage or recombination. Parameter values indicated at the top of the figure. Chromosomes consisting of 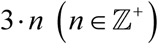 genes are plotted as thick lines.

By allowing chromosome breakage (*ν*_break_ = 0.01; keeping *S* = 30), chromosomes ABC and ABCABC dominate the system (Fig. 2). In this case, shorter chromosomes appear recurrently and, together with chromosomatisation, can form the basis of further adaptability of the system (and avoid the previous “frozen state”, c.f. Fig. 1).

**Fig. 2.**
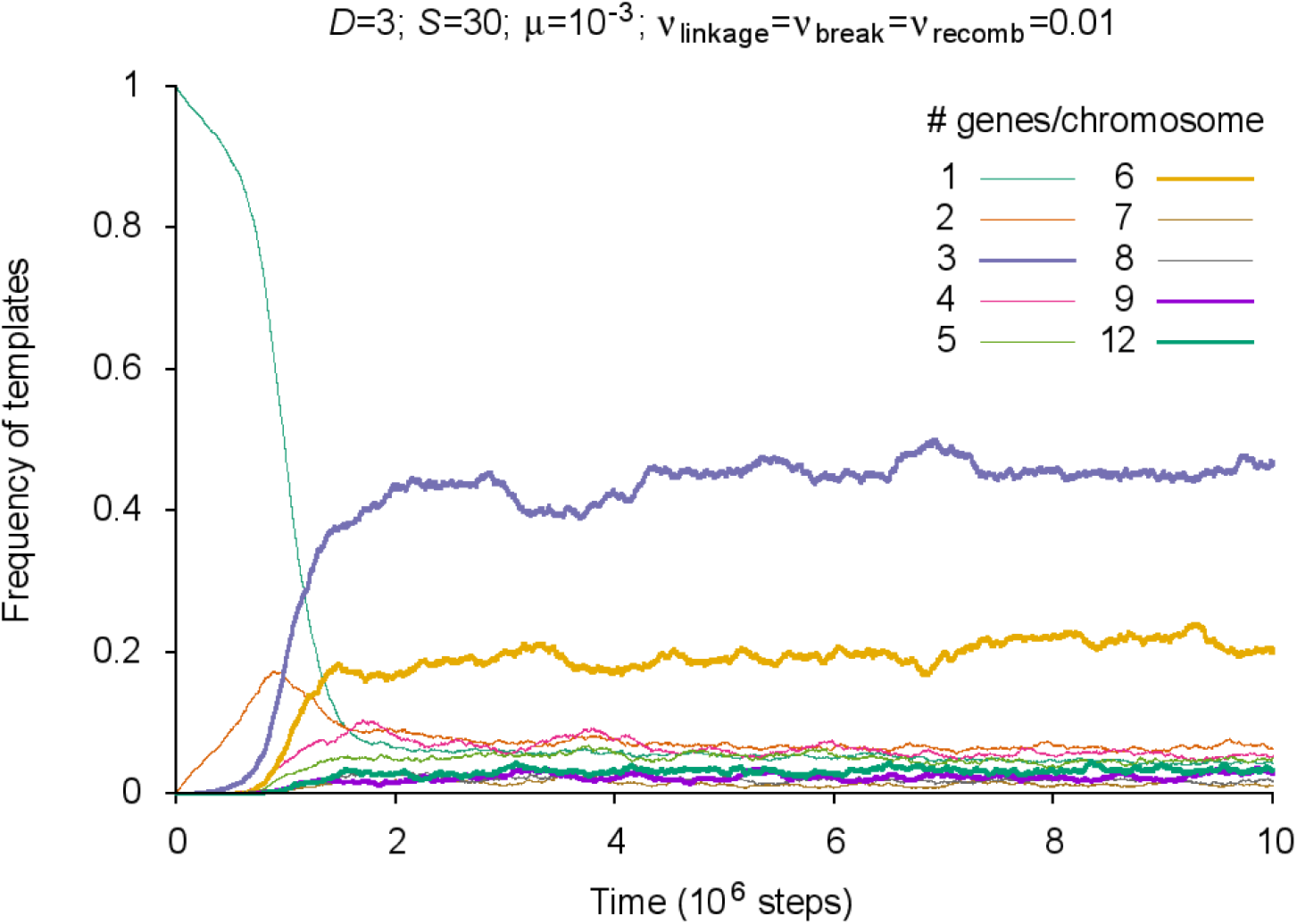
Frequency of different templates normalized on gene count (i.e., a chromosome with 3 genes counts as three when measuring the frequency) with chromosome break and recombination. Parameter values indicated at the top of the figure. Chromosomes consisting of 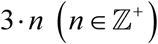 genes are plotted as thick lines (chromosomes with a frequency less than 1% are not shown).

Both split size (*S*) and chromosome breakage (*υ*_break_ > 0) have important effects on the dynamics. If split size is low (*S* = 12), chromosomes with 3 genes dominate because at low split size acquisition of a chromosome with six genes is dangerous due to the early protocell fission (Fig. S2). With higher split size the concentration of chromosomes with six genes (mainly ABCABC type) increases while that of 3-gene chromosomes (mainly ABC type) decreases (Fig. 2). A further increase in split size (*S* = 50) results in decreasing concentration of chromosomes with 3 and 6 genes and increasing number of chromosomes with 9 genes (Fig. S3). High *S* results in a higher amount of longer chromosomes without a full set of genes (Fig. S3). Higher mutation rate does not alter the outcome of the chromosomatization in the sense of the ratio of different type of chromosomes but increases the fluctuation in the frequencies, mainly due to the stochasticity due to the diminished number of viable protocells (Fig. S4).

The higher number of essential genes results in the domination of longer chromosomes with one (or more) full set of genes. Figure S5 shows the result of the simulation with five essential genes (*D* = 5): half of the genes are organized in chromosomes with five genes; the second most frequent is the 10 genes chromosomes class.

Fitness increases with higher split size until the *S* = 20 − 25 region, then slowly drops (Fig. S6). The fitness curve (in the lower split size region, *S* < 25) peaks at *S*\* = *n* ⋅ *D* +1, where *n* ≥ 3, 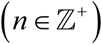. The explanation is clear: if *S* = *n* ⋅ *D*, then the protocell can maintain no more than a *n* − 1 balanced composition of genes before splitting. If, on the other hand, *S* = *n*⋅ *D* +1, then the protocell can maintain one more full set (ABC) of genes (by acquiring a longer chromosome: ABCABCACB instead of the ABCACB type), or harbor instead a new ABC-type chromosome.

Both changes are favorable for protocell’s fitness (a balanced composition of genes at a higher dose) and effectively reduce the assortment load. In the higher *S* region the effect of “proper filling” the protocell with balanced composition right before fission is diminished. With increasing split size the average gene number of chromosomes is increasing, and more or less has the same structure as the fitness curve (at lower *S* region peaks at *S*\* = *n* ⋅ *D* +1 (Fig. S7). With further increase in split size fitness slightly decreases as the strength of group selection weakens (35). A detailed analysis on the effect of different parameters on the outcome can be found in *SI Section 4*.

We have found that the presence/absence of recombination does not affect the quantitative outcomes (frequencies of different types of chromosomes, fitness). (Recombination might help the system to reach the equilibrium state faster, but there is no consistent way to measure this time.) Note that the strong selection on preserving the proper pattern of the recognition site for the replicase (c.f. Eq. (3)) results in one mutated nucleotide (corresponds to 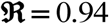) as an average – for a histogram of the number of mutations in equilibrium, see Fig. S8.

### Screening the parameter space

In line with the previous results, the average length of chromosomes increases with both the number of essential genes *D* and the split size *S.* In Fig. 3 one can see the average length of chromosomes (light color: solitary genes, deep blue: chromosomes with 7 genes) as a function of the number of essential genes *D* and the split size *S* assuming *μ* = 10^−3^. Chromosomatization, by decreasing the assortment load, effectively increases the sustainable number of genes: the area enclosed in black lines in Fig. 3 shows the viable region without chromosomatisation. Fig. S9 shows the viable region at different mutation rates. For details, see *SI Section 5*.

**Fig. 3.**
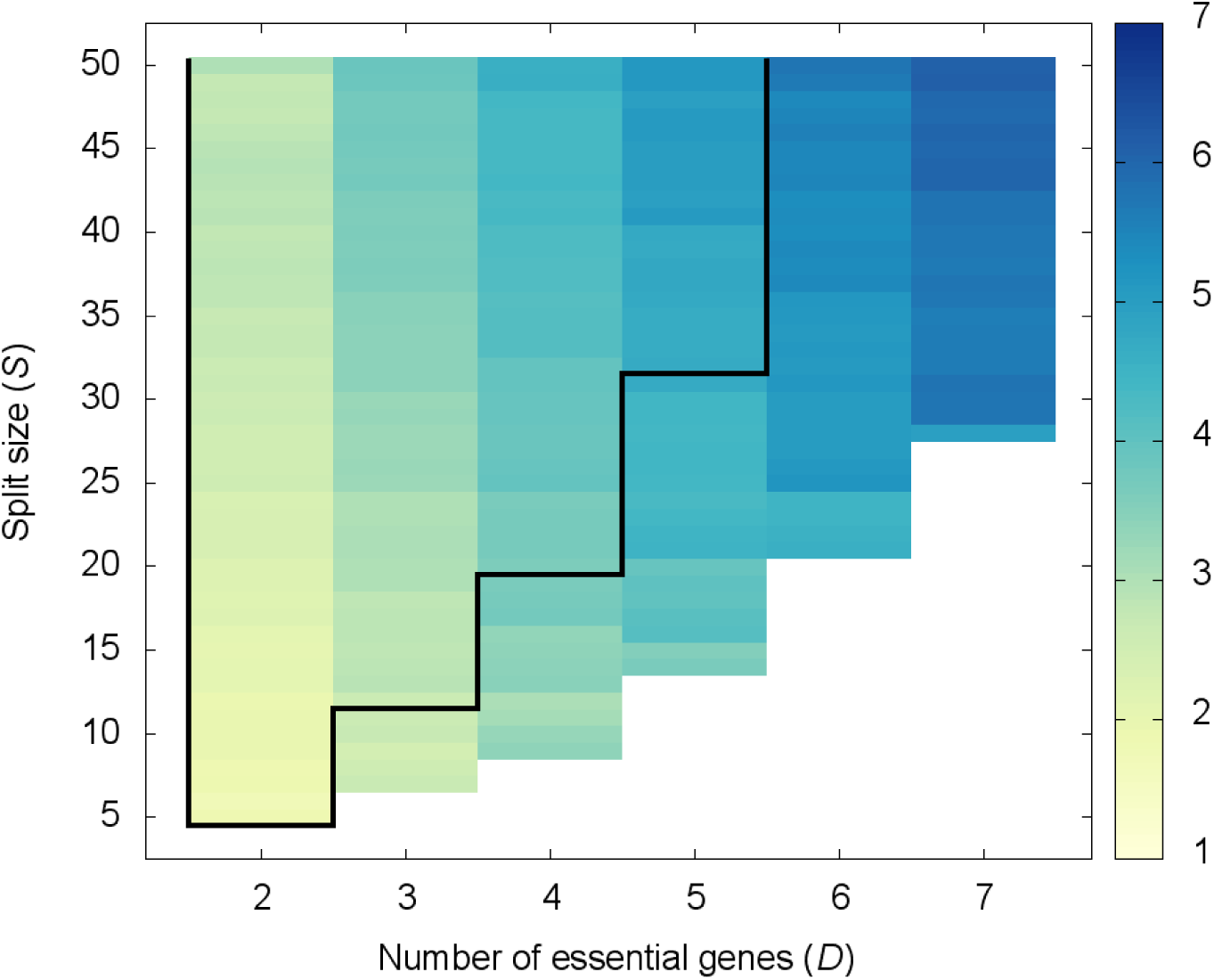
Average number of genes in chromosomes as a function of gene number (*D*) and split size (*S*) with break and recombination. Parameters are *μ*=10^−3^, ν_linkage_ = ν_break_ = ν_recomb_ =0.01. The area enclosed in black lines shows the viable region without chromosomatization.

### The effect of fast replicating parasites

We have analyzed the effect of parasitic genes; that is, genes with high replication rates (higher individual-level selection than metabolic genes) that lack metabolic activity (do not contribute to the collective-level selection).

To check the evolutionary stability of the system against parasites, we changed a given amount of genes to parasites at *t* = 0. Introducing parasites at the beginning is the worst case scenario for the system due to the strong template competition within protocells, and the lack of the beneficial effect of reduced assortment load caused by chromosomes as they are not yet present.

For the whole tested parameter space, we have found that the system is robust against parasites even if their replicative advantage is unrealistically high (50% higher than that of genes with proper recognition site for replicase) and, at the beginning, 25% of templates’ composition within protocells are parasitic molecules. Because of the strong effect of stochastic correction, the frequency of parasites started to decrease in the second generation (after the first replication of all protocells as an average) and basically faded away from the population after approximately 16 generations (Fig. S10). Remarkably, the stochastic correction is so strong that we have not found coexistence between metabolic genes/chromosomes and parasites in the entire investigated parameter space (*SI Section 6*).

## Conclusions

In the “bag of genes” protocell (namely, the stochastic corrector) model (5) chromosomes must make a difference, because they decrease the assortment load (gene A is likely to find its synergistic partner gene B “in the same boat”) and alleviate intragenomic conflict (genes on the same chromosome are necessarily co-replicating (14). This established knowledge suffered from two potential threats: the unknown effects of gene dosage and the mutational load. Our results here have annihilated these threats with promising results. We found that the gene dosage effect selects for balanced gene compositions in emerging chromosomes, and −somewhat counterintuitively − that there is a tendency for the formation of long chromosomes with “multigene families”, also with a dosage-balanced gene composition.

Noteworthy is the fact that the number of sustainable gene types increases with chromosome formation. This we primarily attribute to the considerably decreased assortment load, because the latter increases with the number of gene types without linkage (for the same split size). Remarkably, unlinked genes do not beat chromosomes even for low number of gene types (*D* = 2), very high mutation rates (*μ* = 7 ⋅10^−3^ − 8⋅10^−3^) and high split size. In the modelled context (selection for high metabolic flux, since that ensure fast protocell growth) we find antagonistic epistasis between pairs of gene types, a fact that also favors linkage. Furthermore, the system is remarkably resistant against parasitic mutants. The effect of the emerging multigene families combined with the dosage effect and recurrent mutations warrants detailed analysis of the mutational load that will be presented elsewhere.

A major finding is that the stochastic corrector mechanism prevails, but is shifted from gene to chromosome level. This makes sense because there is yet no accurate segregation mechanism, hence selection favors multiple chromosomes; otherwise the assortment load at the chromosome level would be prohibitive. Thus chromosomes beat genes in the simulated model, but only if the former have sufficiently high copy numbers. The dosage effect selects not only for several gene copies to be maintained, but also for chromosomes harboring balanced multigene families. This genomic composition is expected to disappear with accurate chromosome segregation and efficient transcription of genes in a later stage of evolution (awaiting further work).

Chromosome formation is a critical stage of the first major evolutionary transition (13). It solidifies the protocell level of evolution (“social group maintenance” *sensu* Bourke; see ref. 36). It also enables the appearance of truly specific enzymes, since without linkage inefficient but multifunctional enzymes are selected for (1). Note that here we did not model this aspect in that we assumed that enzymatic functions of genes are efficient and chemically orthogonal. A task for the future is to simulate the coevolution of enzymatic specificity/promiscuity and chromosome formation.

A further unknown is the combined effect of chromosome formation and sex between protocells. The latter is good without chromosomes when protocells with (partial) aneuploidy are more likely to fuse than healthy cells (37). There are two potential levels of mixing, however: the reshuffling of genes and chromosomes between fusing protocells, and molecular recombination among chromosomes. Given antagonistic epistasis the latter is expected to be detrimental, but the complexity of the situation may give us surprises. We believe that the evolutionary origins of a primitive prokaryote-like genome organization will be clarified within the next few years in the context of comprehensive models integrating the discussed features.

## Acknowledgements

The authors are supported by the ATTRACT Project EmLife; by the Wolkswagen Foundation (initiative “Leben? –Ein neuer Blick der Naturwissenschaften auf die grundlegenden Prinzipien des Lebens”, project “A unified model of recombination in life”); by the National Research, Development and Innovation Office (NKFIH) under OTKA grant numbers K124438 (A.S.), K119347 (E.S. and A.S.) and GINOP-2.3.2-15-2016-00057 (E.S. and A.S.) research grants; and by grants CGL2017-89160-P from the Ministerio de Economía, Industria y Competitividad (M.S.) and 2017 SGR 01379 from Generalitat de Catalunya (M.S.). A.S. was supported by the Bolyai János Research Fellowship of the Hungarian Academy of Sciences and the ÚNKP-18-4 New National Excellence Program of the Ministry of Human Capacities. We thank András Hubai and Ádám Kun for stimulating us to revisit the chromosome formation problem and to Tamás Czárán and István Scheuring for comments

## Author contributions

A.S., V.P.K., and M.S. performed computer simulations; A.S. and E.S. contributed the analytical results; A.S., E.S., and M.S. wrote the paper.

The authors declare no conflict of interest.

