## Supplementary Information for "Evolution of linkage and genome expansion in protocells"

### Contents


| | | Page |
|---|---|---|
| 1. | Protocell fitness is maximum if all genes (enzymes) have uniform total activity | 3 |
| 2. | A restricted extremum behavior of the fitness function (compensatory mutations) | 4 |
| 3. | Calculation of epistasis in some simplified cases | 5 |
| | Figure S1 | 6 |
| 4. | Results of different runs | 6 |
| | Figure S2 | 7 |
| | Figure S3 | 8 |
| | Figure S4 | 9 |
| | Figure S5 | 10 |
| | Figure S6 | 11 |
| | Figure S7 | 12 |
| | Figure S8 | 13 |
| 5. | Screening the parameter space | 13 |
| | Figure S9 | 14-15 |
| 6. | The effect of fast replicating parasites | 15 |
| | Figure S10 | 16 |
| References | | 16 |

### 1 Protocell fitness is maximum if all genes (enzymes) have uniform total activity

For simplicity we will use the unnormalized fitness function of Eq. (1) in the main text

$$F = \frac{1}{\sum_{i=1}^{D}\left(\frac{1}{\sum_j g_{ij}A_{ij}}\right)} \tag{S1}$$

and would like to find its maximum. The problem can be simplified by finding the maximum of

$$F = \frac{1}{\sum_{i=1}^{D}\frac{1}{e_i}}, \tag{S2}$$

where $e_i = \sum_j g_{ij}A_{ij}$, a combined measure of copy number and activity (the total, weighted activity of enzyme type $i$ ). The maximum is subject to constant total activity $\sum_i e_i = C.$

As a first step we introduce the Lagrange function

$$L(e_1, e_2, ..., e_D) = \frac{1}{\sum_{i=1}^{D}\frac{1}{e_i}} + \lambda\left(\sum_{i=1}^{D} e_i - C\right), \tag{S3}$$

or in more compact form

$$L(\mathbf{e}) = F(\mathbf{e}) + \lambda\left(\sum_{i=1}^{D} e_i - C\right), \tag{S4}$$

where $\mathbf{e} = (e_1, e_2, ..., e_D)$ is a vector of activities.

In a maximum subjected to the total activity constraint $\frac{\partial L(\mathbf{e})}{\partial e_i} = 0$ (for all $i$ ). After simple calculations we arrive at

$$\frac{\partial L(\mathbf{e})}{\partial e_i} = \frac{F^2(\mathbf{e})}{e_i^2} + \lambda. \tag{S5}$$

From this, in maximum $\hat{e}_i = F(\mathbf{e})\sqrt{-\frac{1}{\lambda}}$ . Substituting back into the total activity constraint, after simple rearrangement we get $\lambda = -\frac{D^2F^2(\mathbf{e})}{C^2}$. Combining this with the condition $\frac{\partial L(\mathbf{e})}{\partial e_j} = 0$ we arrive at $\hat{e}_i = \frac{C}{D}$, which means that the protocell fitness has its maximum if all enzymes have the same total activity.

#### 2 A restricted extremum behavior of the fitness function (compensatory mutations)

Let us assume here that both enzyme activity (*A*) and copy number (*g*) are the same for all types of genes. By introducing $e = Ag$ and using the unnormalized fitness function as above

$$F = \frac{1}{\sum_{i=1}^{D}\left(\frac{1}{\sum_j g_{ij}A_{ij}}\right)} = \frac{1}{\sum_{i=1}^{D}\left(\frac{1}{e}\right)} = \frac{e}{D}. \quad \text{(S6)}$$

We will prove that if for one gene the *e* value (the combination of activity and copy number) reduces by an amount $d(<i)$ and, as a compensatory effect, for one other type increases by *d*, the metabolic flux will decrease.

The calculation of the unaltered flux can also be done in the following way

$$F = \frac{1}{\sum_{i=1}^{D}\frac{1}{e}} = \frac{1}{\binom{D}{D-1}\prod_{i=1}^{D-1}e \Big/ \prod_{i=1}^{D}e} = \frac{\prod_{i=1}^{D}e}{D\prod_{i=1}^{D-1}e} = \frac{e}{D}. \quad \text{(S7)}$$

By introducing the $e \to e+d$ and $e \to e-d$ for any two gene types, the modified fitness can be expressed as

$$F' = \frac{(e+d)(e-d)e^{D-2}}{(D-2)(e+d)(e-d)e^{D-3} + (e+d)e^{D-2} + (e-d)e^{D-2}}. \quad \text{(S8)}$$

After some simple manipulation we get the following result

$$F' = \frac{e}{\left(D + 2\frac{d^2}{e^2+d^2}\right)} < F, \quad \text{(S9)}$$

which indicates that fitness reduces even if the mutation is compensatory.

#### 3 Calculation of epistasis in some simplified cases

*Two genes*. Let us denote the two unmutated gene (enzyme) types by *A* and *B*, and when they carry a deleterious mutation by $A'$ and $B'$. The activity of each unmutated enzyme is 1, and that of the mutated enzyme is $c(<1)$. We assume that both enzymes are present in one copy. From Eq. (1) in the main text, the fitness of different combinations can be easily calculated

$$w_{AB} = 1, \;\; w_{A'B} = w_{AB'} = 2\frac{c}{c+1}, \; w_{A'B'} = c, \quad \text{(S10)}$$

where we assume $D = S = 2$ to get normalized fitness.

Let us define epistasis $(\varepsilon)$ as the fitness decrease of the double mutant divided by twice the fitness decrease of a single mutation

$$\varepsilon = \frac{(1-c)}{2\left(1-2\dfrac{c}{c+1}\right)} = \frac{(1+c)}{2} < 1, \qquad \text{(S11)}$$

i.e., the epistasis is positive. If we assume that both genes are present in $g$ each copies, a similar calculation gives the same result (in this case $S = 2g$ ).

*More genes.* Let us assume that there are $D$ different gene types. The activity of the wild type and the mutant are 1 and $c$, respectively. Let $m$ denote the number of different gene types for which at least one copy is mutant, thus $D - m$ gene types lack mutation. We assume $g$ number of copies for each types of gene. For a mutated gene type $f$ copies bear a single mutation, and $g - f$ copies are wild type. With these assumptions the fitness can be calculated as $(S = Dg)$

$$F = \frac{Dg\left[g - f(1-c)\right]}{D\left[g - f(1-c)\right] + mf(1-c)}. \qquad \text{(S12)}$$

The fitness as a function of $m$ and $f$ is clearly nonlinear, indicating epistatic effects. The strength of the epistasis depends on parameter values. $g - f(1-c)$ is always positive as $f \le g$ and $0 < c < 1$, thus the $F(m)$ function is always convex downward, clearly indicating positive epistasis (on both additive and multiplicative scales; c.f. Fig. S1).

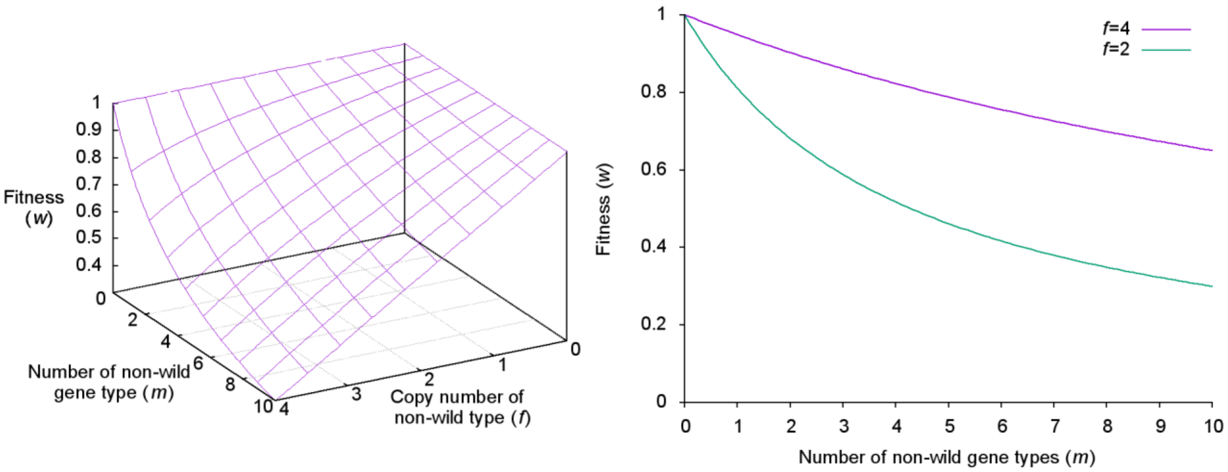


**Fig. S1:** The $F(f, m)$ function (left panel) and the $F(m)$ function at $f = 2$ and $f = 4$ (right panel). Parameters are $D = 10$, $c = 0.3$, $g = 4$.

### 4 Results of different runs

We plot here a number of graphs varying different parameters as indicated in the figures. We compare these simulations to the "reference run" of Fig. 2 ($D = 3$, $S = 30$, $\mu = 10^{-3}$,

$\nu_{\text{linkage}} = \nu_{\text{break}} = \nu_{\text{recomb}} = 0.01$). In the following, we will indicate the changed parameters only. Fig. S2 shows the effect of smaller split size ($S = 12$). The result clearly indicates that smaller split size reduces the amount of larger chromosomes, because acquisition of a chromosome with six genes (ABCABC-type) is dangerous due to the early protocell fission.

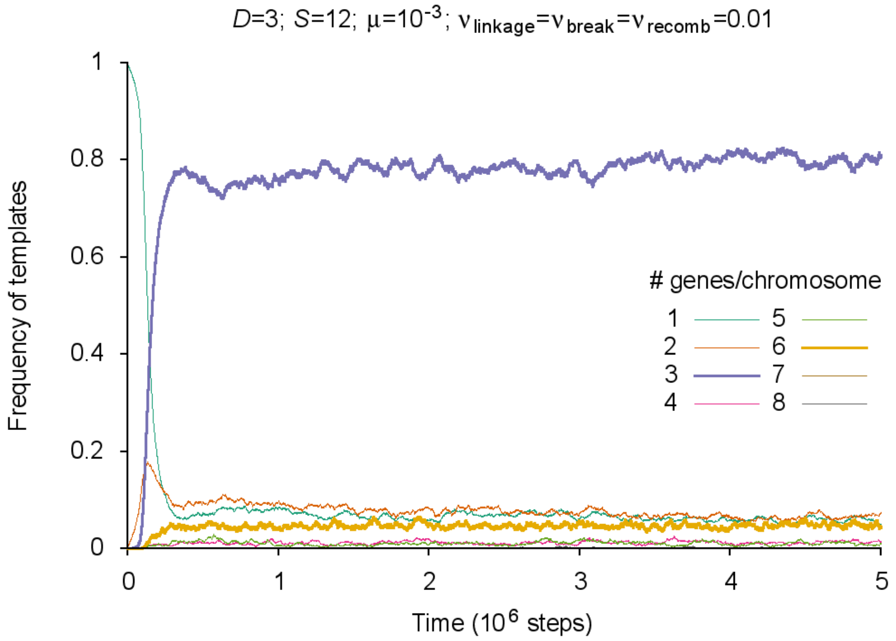


**Fig. S2:** Frequency of different templates normalized on gene count (i.e., a chromosome with 3 genes counts as three when measuring the frequency). Chromosomes consisting of $3 \cdot n \ (n \in \mathbb{Z}^+)$ genes are plotted as thick lines (chromosomes with a frequency less than 2% are not shown).

A simulation presented in Fig. S3 shows the opposite situation; the effect of larger split size $(S = 50)$. This split size allows a higher concentration of chromosomes with 9 genes, as these chromosomes cannot cause too early division. Parallel with the increasing concentration of 9-genes chromosomes, the frequency of 3-genes chromosomes reduces while the frequency of 6-genes chromosomes remains the same. Higher split size results in a higher total amount of no $n \cdot D$ type chromosomes that cannot have one or more full set of essential genes.

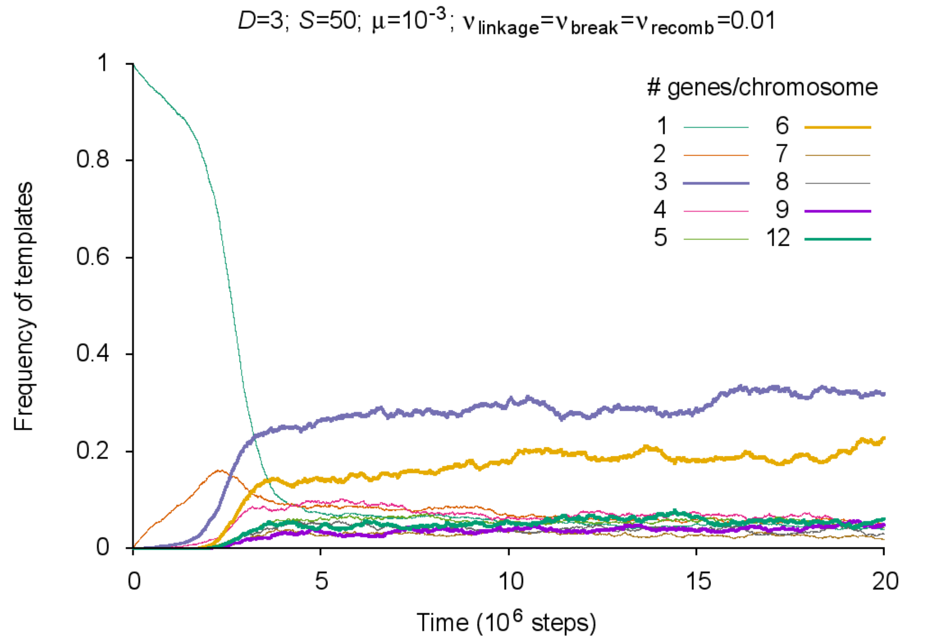


**Fig. S3:** Frequency of different templates normalized on gene count (i.e., a chromosome with 3 genes counts as three when measuring the frequency). Chromosomes consisting of $3 \cdot n \ \left(n \in \mathbb{Z}^+\right)$ genes are plotted as thick lines (chromosomes with a frequency less than 2% are not shown).

We have also investigated the effect of higher mutation rate (Fig. S4). The qualitative behaviour does not change and the ratio of different types of chromosomes remained mainly unaltered (cf. Fig. 2). The fluctuation in the frequency is mainly due to the stochasticity generated by the lower concentration of viable protocells (approximately 35% of the protocells have nonzero fitness, and the mean fitness is about 0.02).

If the number of essential genes is higher longer chromosomes appear. The dominant chromosome class consists of one or more full set of genes (they are of  types). Fig. S5 shows the result of a simulation with $D = 5$, where the 5 genes chromosomes (almost all are ABCDE-type) dominate the system and the second most populated class is the class of 10 genes chromosomes. At $S = 30$, the 15-genes chromosomes cannot appear as these chromosomes cause immediate protocell fission.

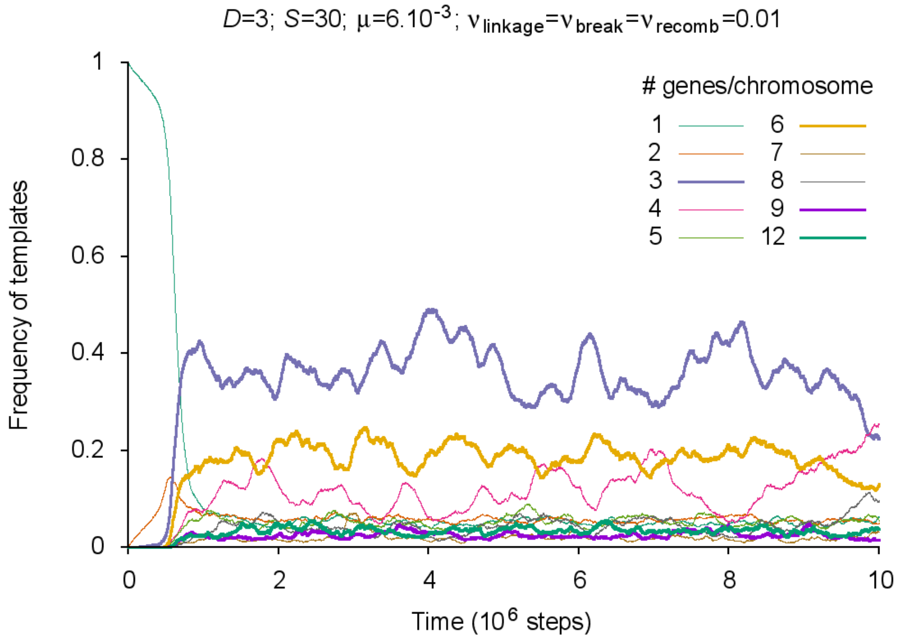


**Fig. S4:** Frequency of different templates normalized on gene count (i.e., a chromosome with 3 genes counts as three when measuring the frequency). Chromosomes consisting of $3 \cdot n$ $\left(n \in \mathbb{Z}^{+}\right)$ genes are plotted as thick lines (chromosomes with a frequency less than 2% are not shown).

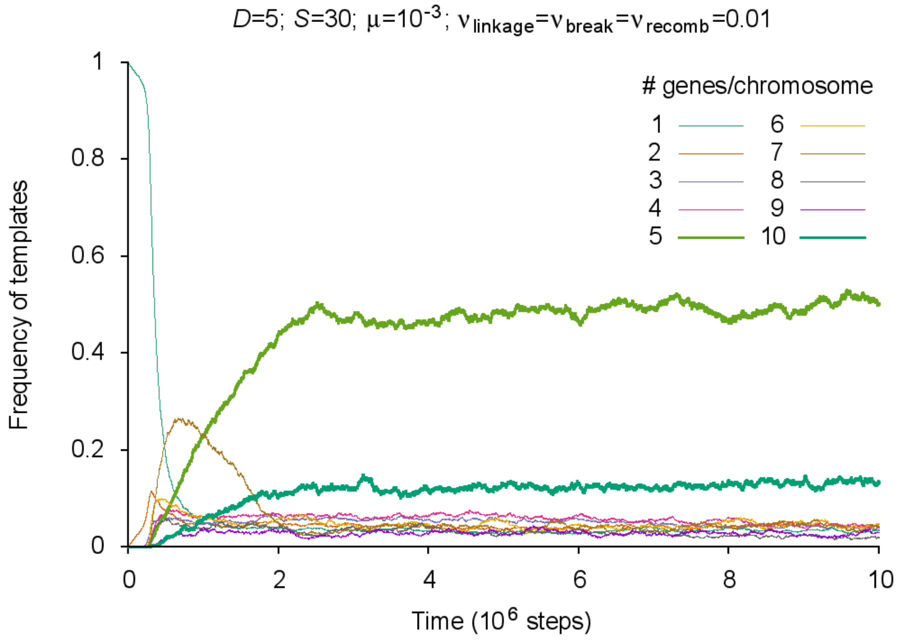


**Fig. S5:** Frequency of different templates normalized on gene count (i.e., a chromosome with 3 genes counts as three when measuring the frequency). Chromosomes consisting of $3 \cdot n$ $\left(n \in \mathbb{Z}^{+}\right)$ genes are plotted as thick lines (chromosomes with a frequency less than 2% are not shown).

We have also investigated the fitness and the average gene number of chromosomes as a function of the split size (*S*). As it is known from the theory of group selection [1], if the size of the group is too large the fitness tends to drop. Figure S6 clearly indicates this behavior. The optimal group size for the parameters indicated in Fig. 2 is about 20–25. The fitness curve (in the lower split size region, $S < 25$) peaks at $S^* = n \cdot D + 1$, where $n \geq 3, \left(n \in \mathbb{Z}^+\right)$, because if $S = n \cdot D + 1$ then the protocell can maintain one more full sets of genes by acquiring a longer chromosome. For detailed explanation of the structure of the fitness curve, see the main text.

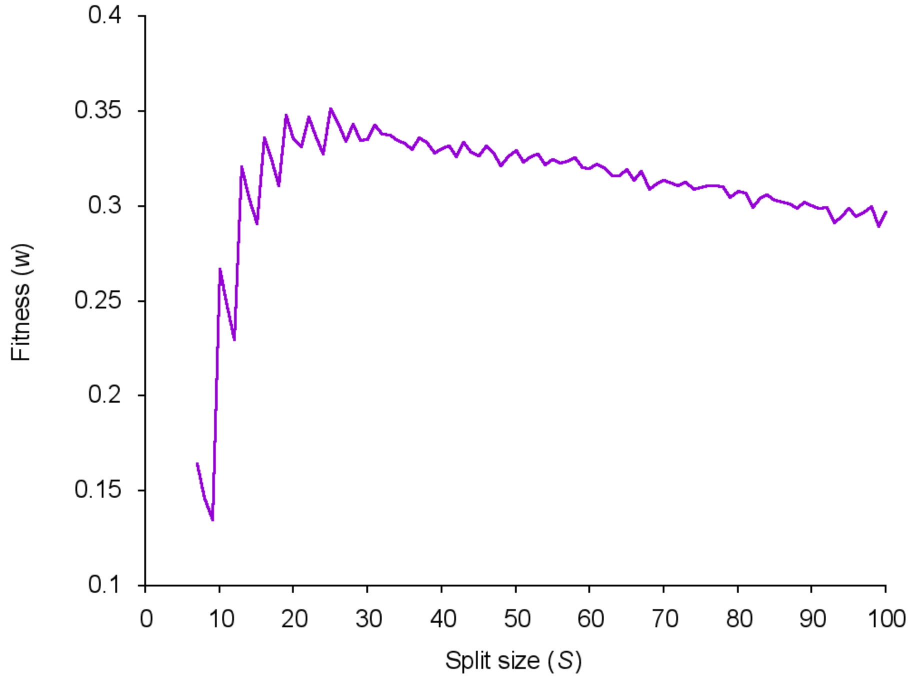


**Fig. S6:** The equilibrium fitness of the population as a function of the split size (*S*). Average of 10 independent runs. Relevant parameters as in Fig. 2 $\left(\mu = 10^{-3};\ \nu_{\text{linkage}} = \nu_{\text{break}} = \nu_{\text{recomb}} = 0.01\right)$.

The course of the average gene number of chromosomes as a function of the split size (Fig. S7) has a saturating characteristic. In the $10 < S < 60$ region of split size the gene number increases with the split size in a linear way, indicating the dosage effect: more balanced composition of genes results in higher fitness. This effect diminishes if $S \gg D$; cf, the $S > 70$ region in Fig. S7. With higher number of essential genes (*D*) the beginning of the saturated regime becomes at higher split density (data not shown).

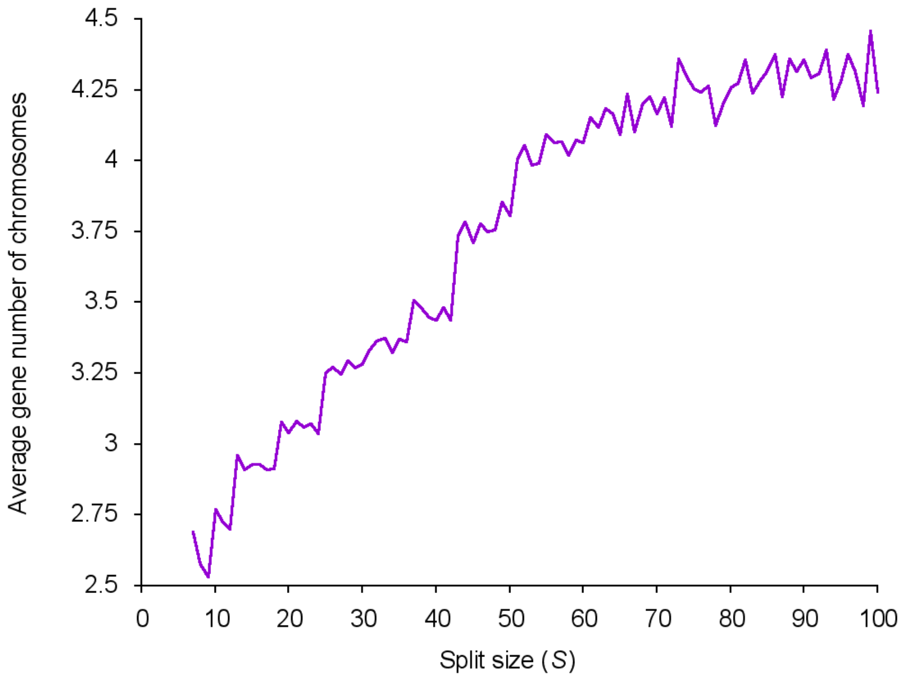


**Fig. S7:** Average gene number of chromosomes (averaged over the population) as a function of the split size. Average of 10 independent runs. Relevant parameters as in Fig. 2 $\left(\mu = 10^{-3};\ \nu_{\mathrm{linkage}} = \nu_{\mathrm{break}} = \nu_{\mathrm{recomb}} = 0.01\right)$

Fig. S8 shows the distribution of the number of mutations in the region defining the target affinity towards the replicase in the evolved population (at $t = 10^7$). The peak at one mutated nucleotides corresponds to $\mathfrak{R} = 0.938$ replication probability, cf. Eq. (3) in the main text. As the replication probability $\mathfrak{R}$ is a fast decreasing function of the number of mutated nucleotides $\psi_t$, the replication probability of genes/chromosomes with more than four errors in the relevant region is negligible.

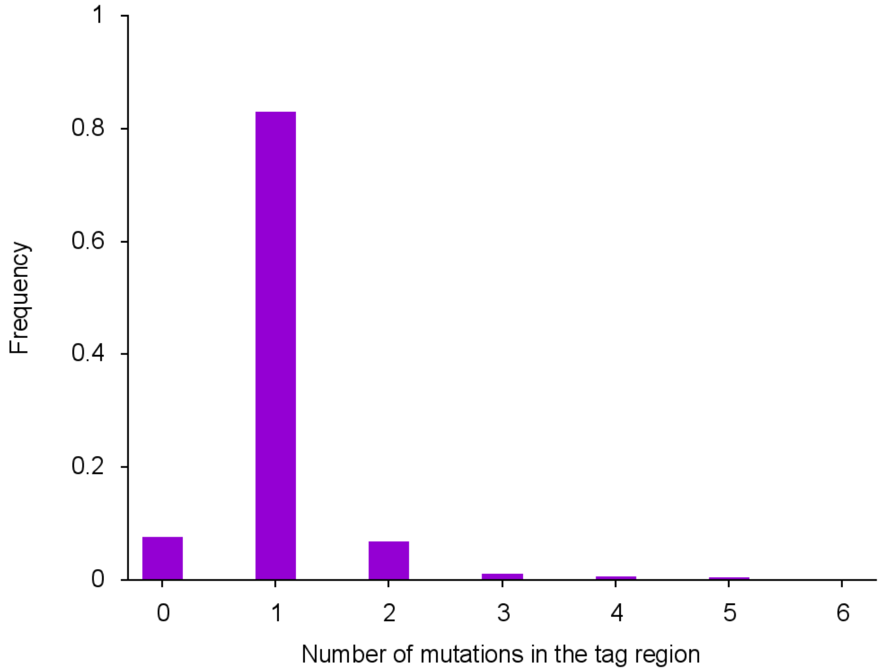


**Fig. S8:** Distribution of the number of mutated nucleotides in the region defining the target affinity towards the replicase. Parameters are the same than in Fig. 2. The values are averaged over 25.000 time steps at $t = 10^7$. Note that, according to Eq. (3) in the main text, the affinities corresponding to 0, 1, 2 and 3 mutated nucleotides are $\Re = 1;\ 0.938;\ 0.319;\ 0.058$, respectively.

#### 5. Screening the parameter space

We have also analyzed the average gene number of chromosomes in the parameter space spanned by the split size (*S*), the number of essential genes (*D*), and the mutation rate (μ). Different panels of Fig. S9 correspond to different mutation rates from $\mu = 0$ to $\mu = 8 \times 10^{-3}$ (increasing mutation rates from left to right and from top to bottom). Each plot shows the average gene number (with color coding) as an average over 200,000 time steps starting at $t = 2 \times 10^6$. The periodic pattern of the average number of genes at fixed $D$ is in agreement with the observation of the special role of $S^* = n \cdot D + 1$ split size, at which split density the average gene number increases as one more full set of genes can be harbored (see Results and Discussion in the main text). The non-coherent fine structure at higher *D* (and higher *S*) regime is mainly due to the stochastic effects. It can be seen that the viable region in the *S*-*D* planes shrinks: with the increasing mutation rate the system can maintain fewer types of genes.

The effect of chromosomatization on the sustainable amount of information (the number of sustainable essential genes, *D*) has also been investigated. We ran simulations with no chromosomatization $\left(\nu_{\text{linkage}} = \nu_{\text{break}} = \nu_{\text{recomb}} = 0\right)$ and the viable region of the parameter space enclosed in black lines. The possibility of chromosome formation increased the number of sustainable genes by a factor of 2 to 3.

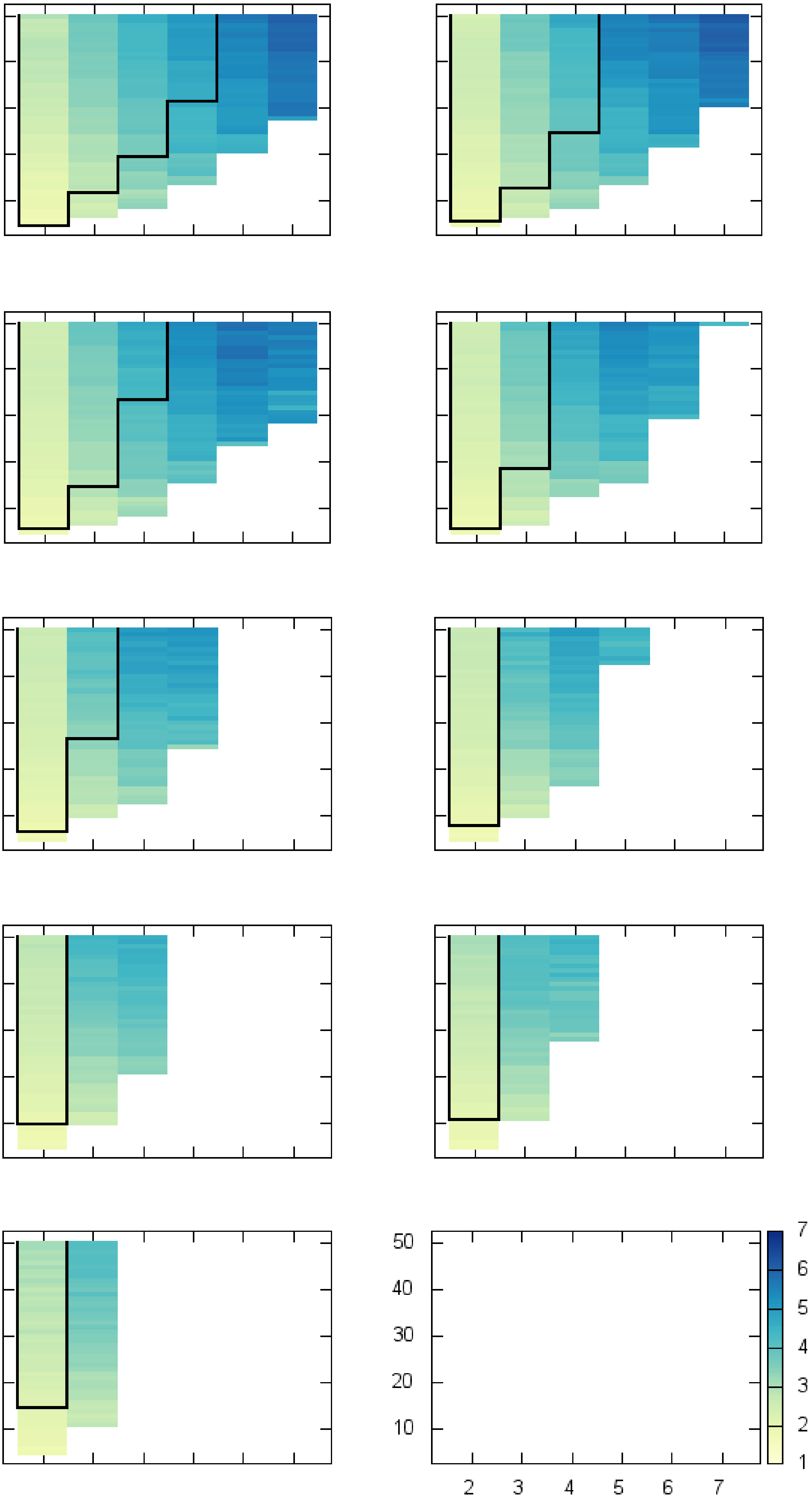
50
40
30
20
10
2
3
4
5
6
7
1
2
3
4
5
6
7

**Fig. S9:** Average number of genes in chromosomes (color bar) as a function of gene number ($D$, $x$-axis) and split size ($S$, $y$-axis), with breakage and recombination at different mutation rates (from left to right and top to bottom: $\mu = 0, 10^{-3}, 2\times10^{-3}, 3\times10^{-3}, \cdots, 8\times10^{-3}$). Parameters are: $\nu_{\text{linkage}}=\nu_{\text{break}}=\nu_{\text{recomb}}=0.01$. The area enclosed in black lines shows the viable region without chromosomatization.

#### 6. The effect of fast replicating parasites

We have analyzed the effect of parasites; that is, genes with higher $\mathfrak{R}$ affinity towards the replicase and without metabolic activity. We have made a series of runs over the relevant part of parameter space $(D, S, \mu, \nu)$ with different replicase affinities $\mathfrak{R}$ and different concentrations of parasites. In order to analyze the worst case scenario for the system, we made two assumptions: i) we add parasites at $t$ = 0 and, thus, parasites compete mainly with singe genes to ignore the reduced assortment load caused by chromosomes; and ii) we assumed that mutations do not act on the region that defines parasites' activity towards replicase (i.e., constant affinity towards the replicase). Linkage between parasites was ignored. We have found that in the entire part of the parameter space used in the simulations, the parasites disappear from the system. Remarkably, stochastic correction is so strong that we have not found coexistence between metabolic genes/chromosomes and parasites.

Fig. S10 shows the time course of the frequency of parasites with unrealistically high replication rate $\mathfrak{R}=1.5$ (50% higher than the maximum for non-parasites) and a very high initial concentration (25% of the genes are parasites). As one can see, the stochastic correction eliminates parasites relatively fast. The characteristic time of a generation (the time necessary for division of all vesicles at an average) can be estimated as $\tau = \frac{S}{2}N$ , where $S$ is the split size and $N$ is the number of protocells in the population. With the parameters of Fig. S10 $\tau = 75{,}000$; exclusion of the parasite starts during the second generation and finishes around the 16th generation.

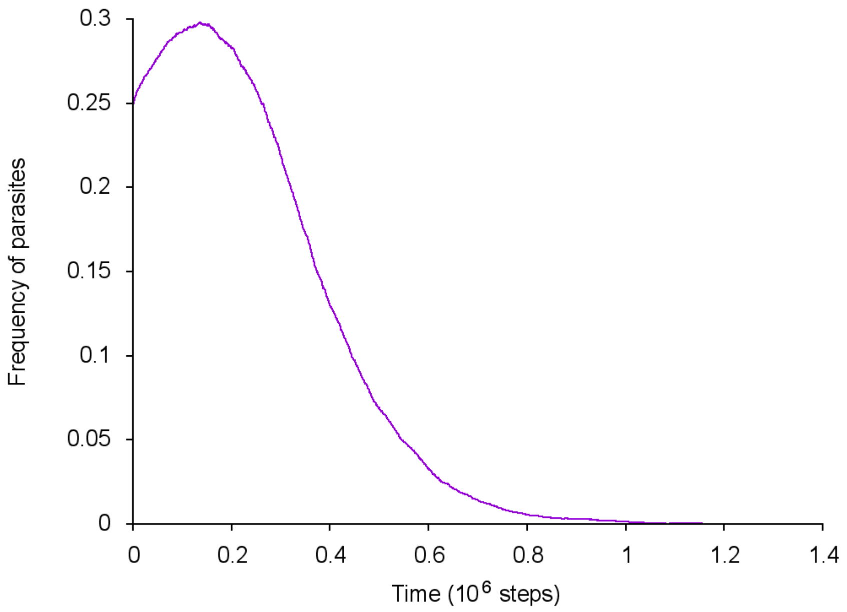


**Fig. S10:** Time course of the frequency of parasites. The affinity of the parasites toward replicase is $\mathfrak{R}=1.5$, and their frequency at $t$ = 0 is 0.25. Other parameters are as in Fig. 2 $(\mu = 10^{-3};\ \nu_{\text{linkage}} = \nu_{\text{break}} = \nu_{\text{recomb}} = 0.01)$.
